# Tracing the environmental footprint of the *Burkholderia pseudomallei* lipopolysaccharide genotypes in the tropical “Top End” of the Northern Territory, Australia

**DOI:** 10.1101/603886

**Authors:** Jessica R. Webb, Audrey Rachlin, Vanessa Rigas, Derek S. Sarovich, Erin P. Price, Mirjam Kaestli, Linda M. Ward, Mark Mayo, Bart J. Currie

## Abstract

The Tier 1 select agent *Burkholderia pseudomallei* is an environmental bacterium that causes melioidosis, a high mortality disease. Variably present genetic markers used to elucidate strain origin, relatedness and virulence in *B. pseudomallei* include the *Burkholderia* intracellular motility factor A (*bimA*) and filamentous hemagglutinin 3 (*fhaB3*) gene variants. Three lipopolysaccharide (LPS) O-antigen types in *B. pseudomallei* have been described, which vary in proportion between Australian and Asian isolates. However, it remains unknown if these LPS types can be used as genetic markers for geospatial analysis within a contiguous melioidosis-endemic region. Using a combination of whole-genome sequencing (WGS), statistical analysis and geographical mapping, we examined if the LPS types can be used as geographical markers in the Northern Territory, Australia. The clinical isolates revealed that LPS A prevalence was highest in the Darwin and surrounds (n = 660; 96% being LPS A and 4% LPS B) and LPS B in the Katherine and Katherine remote and East Arnhem regions (n = 79; 60% being LPS A and 40% LPS B). Bivariate logistics regression of 999 clinical *B. pseudomallei* isolates revealed that the odds of getting a clinical isolate with LPS B was highest in East Arnhem in comparison to Darwin and surrounds (OR 19.5, 95% CI 9.1 – 42.0; *p*<0.001). This geospatial correlation was subsequently confirmed by geographically mapping the LPS type from 340 environmental Top End strains. We also found that in the Top End, the minority *bimA* genotype *bimA*_*Bm*_ has a similar remote region geographical footprint to that of LPS B. In addition, correlation of LPS type with multi-locus sequence typing (MLST) was strong, and where multiple LPS types were identified within a single sequence type, WGS confirmed homoplasy of the MLST loci. The clinical, sero-diagnostic and vaccine implications of geographically-based *B. pseudomallei* LPS types, and their relationships to regional and global dispersal of melioidosis, require global collaborations with further analysis of larger clinically and geospatially-linked datasets.

**Author Summary:** *Burkholderia pseudomallei* is a pathogenic soil bacterium that causes the disease melioidosis, which occurs in many tropical regions globally and in recent years has emerged in non-tropical regions. Melioidosis has been predicted to affect 165,000 people every year resulting in an estimated 89,000 deaths. Person to person transmission is rare with most cases linked to exposure to the bacterium from the environment. The genetic background of *B. pseudomallei* has been well studied and variably present genes have been linked to distinct melioidosis disease states and geographic regions, however we still need a stronger understanding of the association of genes with geography. Three lipopolysaccharide types exist in *B. pseudomallei* and the prevalence of the lipopolysaccharide genes vary between melioidosis endemic regions, but it is unknown if the lipopolysaccharide genes can be used as geographical markers in a single melioidosis-endemic region. In this study, we used a combination of whole-genome sequencing, statistics and geographical mapping to elucidate if the three lipopolysaccharide genes can be used as geographical markers within the Northern Territory, Australia. We show that the three LPS types have distinct but overlapping geographical footprints within a single melioidosis region and can be used as geographic markers alongside a number of other important variably present *B. pseudomallei* genes.

## Introduction

*Burkholderia pseudomallei* is the causative agent of melioidosis, an often fatal disease that is endemic in tropical regions globally, including the “Top End” of the Northern Territory, Australia (1). Clinical presentations of melioidosis are highly varied (2-5) with pneumonia being the most common presentation (6). Melioidosis mortality rates range from <10% in the NT to >40% in regions of Asia and Southeast Asia where prompt diagnosis and accessibility to drugs and hospital care can be limited (2, 6, 7). Global melioidosis mortality has been predicted to be considerably higher than mortality from dengue and leptospirosis combined and similar to that from measles (8). In addition, the bio-threat status of *B. pseudomallei* and the lack of a current vaccine makes this organism of high importance to public health in many regions globally.

Variable genetic markers in *B. pseudomallei* with known geographical associations include the mutually exclusive *Burkholderia* intracellular motility factor A (BimA) variants *bimA*_Bm_ and *bimA*_Bp_, the filamentous hemagglutinin 3 (*fhaB3*) gene and the *Burkholderia thailandensis*-like flagellum and chemotaxis (BTFC/YLF) gene clusters (9, 10). These genetic markers have been used to elucidate strain relatedness, origin and virulence potential, with *bimA*_Bm_ being strongly linked to neurological melioidosis in Australian studies (9). The lipopolysaccharide (LPS) of *B. pseudomallei*, a type II O-polysaccharide (11), consists of A, B and B2 variants (12). As with *bimA*_Bm_/*bimA*_Bp_, YLF/BTFC, and *fhaB3,* the distribution of LPS genotypes varies between Australia and Southeast Asia (9, 12). In Thailand and Australia, LPS type A is the most abundant, followed by LPS B, whereas LPS B2 has not been detected in Thailand but has been detected in Australia (12). Despite regional differences in LPS genotype prevalence, the distribution of these types across different geographical regions within a single melioidosis-endemic region has not been thoroughly investigated. Elucidating the geospatial patterns of LPS genotypes is potentially important for *B. pseudomallei* virulence studies, the development of LPS-based sero-diagnostics, and for consideration of vaccines that have an LPS component.

In this study, we used a combination of WGS, bivariate logistic regression (with the cluster feature) and geographical mapping to examine the geographical distribution of LPS types in the Top End region. We first investigated LPS geographic correlations using a large number of clinical isolates (n = 1005) from our 28-year Darwin Prospective Melioidosis study (DPMS). We then investigated LPS type distributions for 340 environmental strains collected from known Top End locations to determine whether LPS genotypes for clinical isolates, which have usually only presumptive infection locale data, matched that of the environmental isolates, where location is definite. Lastly, we determined *bimA*_Bm_/*bimA*_Bp_ diversity in the 340 environmental strains to determine whether the *bimA* variants and LPS types share similar geographical footprints in the NT.

## Methods

### Ethics statement

This study was approved by the Human Research Ethics Committee of the NT Department of Health and the Menzies School of Health Research (HREC 02/38). In the present study all data that was analyzed were anonymized.

### *B. pseudomallei* isolates used in this study

A total of 1,345 *B. pseudomallei* isolates from the Top End region were used in this study for bivariate analysis and geographical mapping. 1,005 clinical isolates were included in this study and 999 (excluding the 6 B2 strains) were used for bivariate statistical analysis, and environmental isolates (n = 340) were used for geographical mapping. The 1,005 clinical isolates were primary *B. pseudomallei* isolates corresponding to 1,005 patients enrolled in the 28-year DPMS. The clinical isolates were assigned to one of four Top End geographic regions based on the presumptive location of infection and these regions have been segregated and assigned as outlined by the Australian Bureau of Statistics (https://www.abs.gov.au/; statistical area level 3): Darwin and surrounds; Darwin remote (includes West Arnhem), Katherine and Katherine remote, and East Arnhem (Fig 1). Melioidosis is very uncommon in the southern half of the NT and this region was not included in our study.

**Fig 1.**
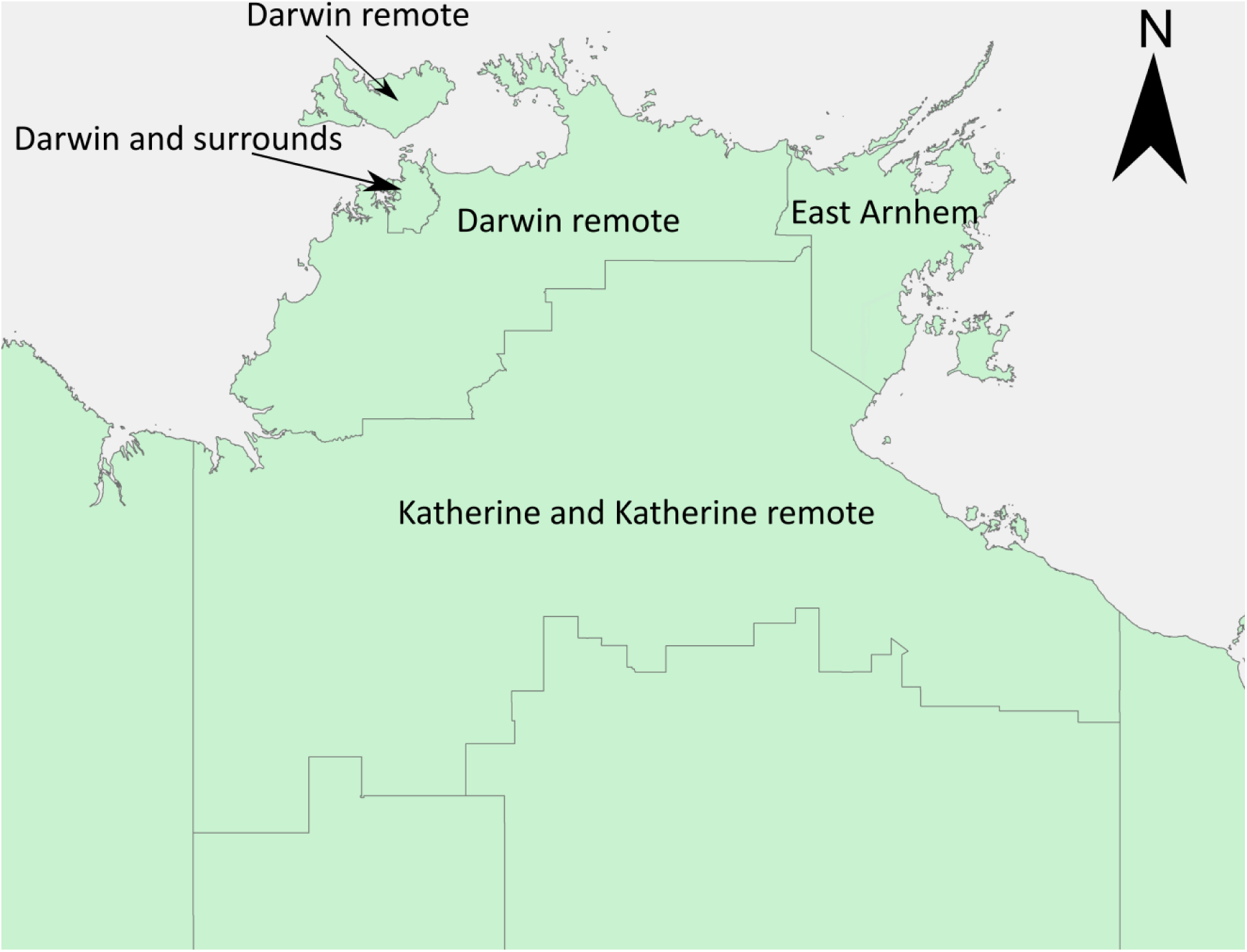
Geographical origin of the *Burkholderia pseudomallei* clinical and environmental isolates within the Top End region of the Northern Territory, Australia, that were used in this study. (adapted from an Australian Bureau of Statistics shape file, of statistical area level 3 (https://www.abs.gov.au/)).

The 340 Top End environmental isolates were processed and subsequently *B. pseudomallei* was confirmed as previously described (13-17). The location for each of the 340 environmental isolates was determined using Global Positioning System coordinates, with mapping of each isolate coordinate using ArcGIS (https://www.arcgis.com/index.html). For comparison we included an additional 61 isolates of Queensland (n = 38), Western Australian (n = 12) and Papua New Guinean (n = 11) origin.

### LPS and *bimA* genotypes

The assignment of LPS types for the 1,005 clinical genomes has previously been determined (18) using the Basic Local Alignment Search Tool (BLAST). The clinical genomes were not assigned a *bimA* variant as this has previously been performed on our clinical strains (10). LPS and *bimA* genotypes were determined for each of the 340 Top End environmental genomes, with an LPS type being assigned to those additional 61 isolates from Western Australia, Papua New Guinea and Queensland, Australia. In brief LPS A (*wbil* to *apaH* in K96243 [GenBank ref: NC_006350]), LPS B (*BUC_3392* to *apaH* in *B. pseudomallei* 579 [GenBank ref: NZ_ACCE01000003]), LPS B2 (*BURP840_LPSb01* to *BURP840_LPSb21* in *B. pseudomallei* MSHR840 [GenBank ref: GU574442]), *bimA*_*Bm*_ (*BURPS668_A2118* in *B. pseudomallei* MSHR668 [GenBank ref: NZ_CP009545]) and *bimA*_*Bp*_ (*BPSS1492* in *B. pseudomallei* K96243) were BLAST-searched against an in-house *B. pseudomallei* database containing all genomes in this study using the nucleotide BLAST (BLASTn) parameter.

### Statistical analysis of LPS type, locale, and multilocus sequence type (MLST) using the clinical *B. pseudomallei* dataset (n = 999)

For each clinical strain, the presumptive geographical location of infection and its sequence type (ST) (https://pubmlst.org/bpseudomallei/) were analysed using Stata version 14.2 (StataCorp LP, College Station, TX, USA). Bivariate logistics regression using the cluster feature was performed for LPS type with locale using the Darwin and surrounding region as the reference level, and bivariate Pearson’s χ^2^ (frequency >5 in all cells) was performed for LPS type with ST or BimA variants: A *P* < 0.05 was considered significant.

### Phylogenomic analysis of 175 *B. pseudomallei* genomes

We performed comparative analyses of 175 *B. pseudomallei* genomes that represent both the global diversity of this bacterium and within-Northern Territory population diversity (Table S1). Of these 175 genomes, 144 were publicly available and 31 were sequenced as part of this study. Genomic DNA for 31 isolates lacking WGS data was extracted as previously described (19). These isolates were sequenced at Macrogen, Inc. (Gasan-dong, Seoul, Republic of Korea) or Australian Genome Research Facility Ltd. (Melbourne, Australia) using the Illumina HiSeq2000 and Illumina HiSeq2500 platforms (Illumina, Inc., San Diego, CA) and these data are available on NCBI (Table S1)

Comparative analysis of the 175 genomes was carried out to investigate geographical clustering of the LPS types, to determine the genetic relatedness of suspected ST homoplasy cases (20, 21), and to ascertain if suspected ST homoplasy cases were intercontinental or intracontinental. Orthologous biallelic single-nucleotide polymorphisms (SNPs) were identified from the WGS data using the default settings of SPANDx v3.2 (22). The closed Australian *B. pseudomallei* genome MSHR1153 (23) was used as the reference for read mapping. A maximum-parsimony (MP) phylogenetic tree was reconstructed based on 219,075 SNPs identified among the 175 genomes using PAUP* 4.0.b5 (24). Recombinogenic SNPs were identified using Gubbins v2.2.0 (default parameters) (25). STs were determined using the BIGSdb tool (26).

## Results

### LPS B is linked to remote Top End regions of the Northern territory

BLAST analysis demonstrated that overall in the Top End, LPS A was dominant in both environmental *B. pseudomallei* strains (A = 89%; LPS B = 10% and LPS B2 = 1%) and in clinical strains (LPS A = 87%; LPS B = 12% and LPS B2 = 1%) (Table 1). Using the clinical *B. pseudomallei* dataset, we performed bivariate logistics regression (with the cluster feature) of LPS type A and B correlations and presumptive infecting locale, using Darwin urban as the reference level. Low LPS B2 numbers (n=6) precluded statistical analysis of this genotype, leaving 999 DPMS isolates for the analysis. Of these six B2 isolates, five were from presumptive infections occurring in Katherine and Katherine remote communities, with the remaining LPS B2 strain linked to infection in the Darwin and surrounding region. Bivariate logistics regression adjusted with the cluster feature demonstrated that the odds of getting a clinical isolate with LPS B in the Darwin remote, Katherine and Katherine remote and East Arnhem regions, respectively is 4.7 (95% CI 2.4 – 9.6; *p*<0.001), 14.6 (95% CI 6.1 – 34.7; *p*<0.001) and 19.5 (95% CI 9.1 – 42.0; *p*<0.001) times higher compared to the odds in the Darwin and surrounding region. LPS B prevalence was highest in patients infected in the Katherine and Katherine remote and East Arnhem regions (n = 79; 64% of LPS B isolates), whereas LPS A prevalence was highest in patients infected in the Darwin and surrounds and Darwin remote regions (n = 756; 86% of LPS A isolates).

**Table 1.**
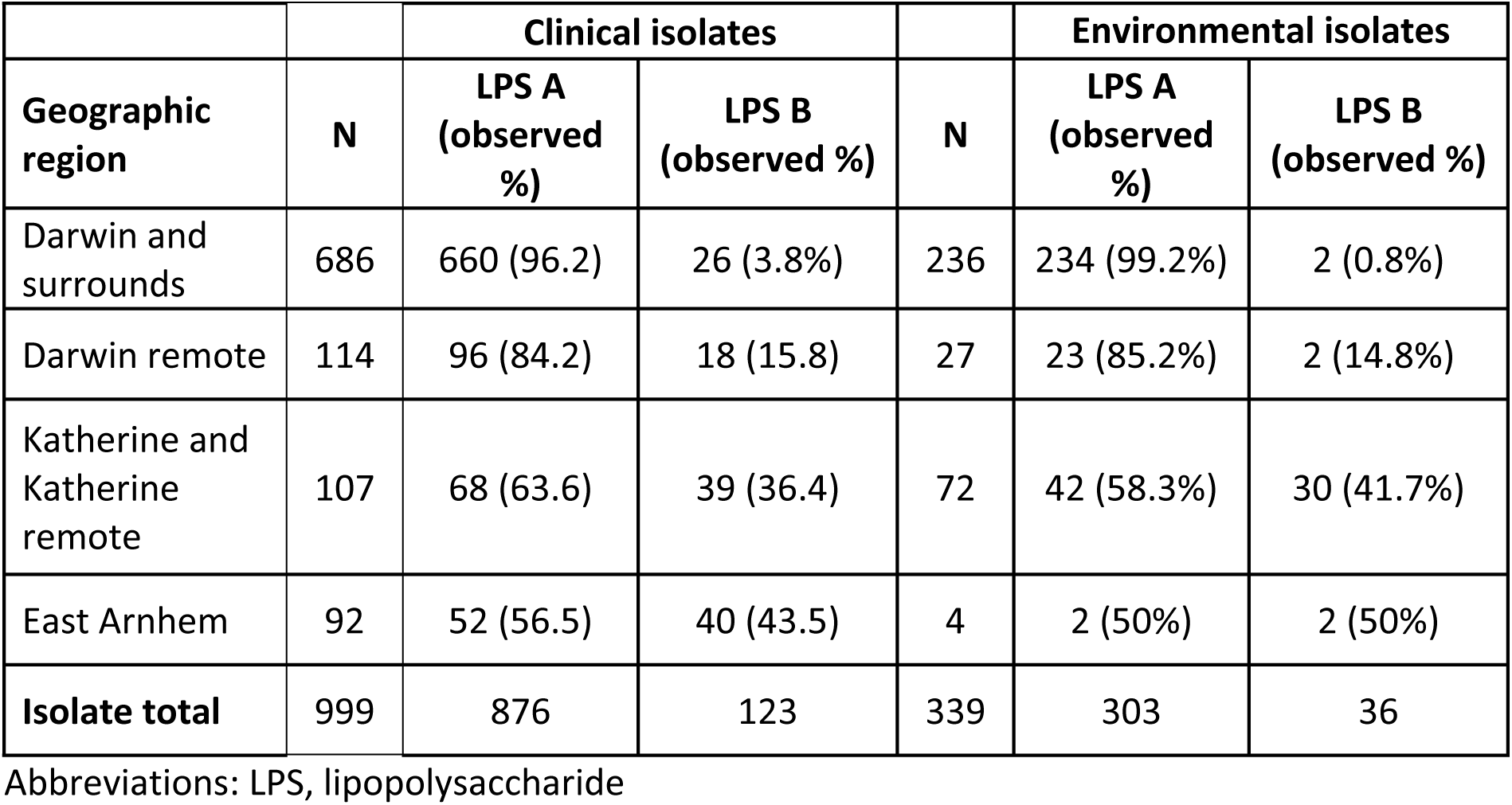
The geographic locale of LPS genotype A and B in *Burkholderia pseudomallei* isolated from melioidosis patients and from the environment in the Top End region, Northern Territory, Australia.

### Geographic clustering of LPS types was also observed in NT environmental strains

Geographical mapping of LPS types for the environmental isolates also supported the link between LPS and geography. LPS type A was predominant in Darwin and surrounds (Table 1 and Fig 2B), whilst LPS B was found in all four geographical regions but with numbers highest in the Katherine and Katherine remote region (Fig 2A and B, number of LPS B strains at each site is indicated). LPS B2 from environmental strains was only observed in the Katherine remote region (n = 1). In non-Top End comparator strains, LPS A was also the dominant genotype (range=63-91%), followed by B (range=9-24%) and B2 (range=0-18%). Numbers of the LPS type A, B, and B2, respectively, were as follows: Western Australia (10, 2, 0), Queensland (24, 9, 5) and Papua New Guinea (8, 1, 2).

**Fig 2.**
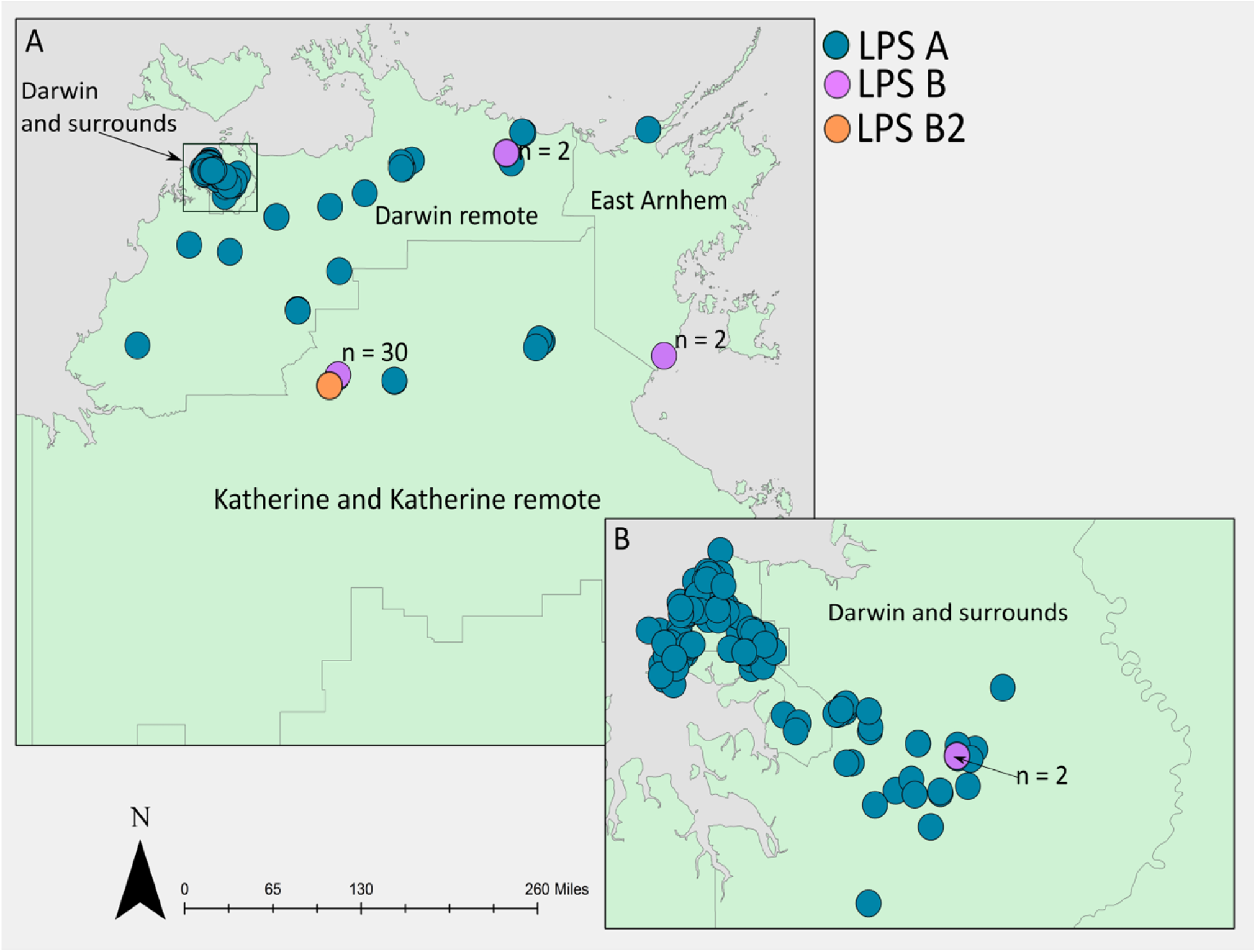
Geographical distribution of LPS genotypes in 340 environmental *Burkholderia pseudomallei* isolates from (A) the Top End region, Northern Territory, Australia and (B) Darwin and surrounds. The number of LPS B strains at each of the four LPS B regions is indicated. ArcGIS (https://www.arcgis.com/index.html) was used for mapping isolates onto an Australian Bureau of Statistics shape file, of statistical area level 3 (https://www.abs.gov.au/).

### Bivariate analysis revealed a strong correlation between LPS A and STs found only in the Darwin and surrounds and Darwin remote regions

Bivariate analysis of clinical isolates encoding LPS types A and B revealed statistically significant associations between LPS types and STs. LPS A was significantly associated with STs only found in the Darwin and surrounding region, including ST-109 (n = 126; 14% of LPS A’s; P<0.001), ST-36 (n = 81; 9% of LPS A’s; P<0.001) and ST-132 (n= 67; 8% of LPS A’s; P<0.001), whilst LPS B was associated with rare STs that are uncommon in the Darwin regions (data not shown). These associations therefore reflected the LPS geographical associations observed in clinical strains. Although LPS B2 was not included in the statistical analysis it was found in strains belonging to rare STs (ST-331 n = 1; ST-737 n = 2; ST-770 n = 2; ST-118 n = 1) with four of the STs unique to the Katherine and Katherine remote region. Of the STs identified in the 1,005 DPMS clinical strains, four contained multiple LPS types amongst 18 strains. Strains belonging to ST-468, ST-807 and ST-734 contained LPS A or LPS B, whilst strains belonging to ST-118 contained LPS A or LPS B2. The LPS and ST discrepancy was also observed in the Top End environmental strain cohort, with three strains belonging to ST-1485 containing LPS A or B.

### The geographical footprint of *bimA*_Bm_ in environmental isolates is similar to LPS B

BLAST analysis demonstrated that *bimA*_Bp_ was the dominant BimA variant in Top End environmental samples (94.8%), similar to the *bimA*_Bp_ prevalence in clinical isolates from a previous study, whereby 95.7% of clinical isolates from Darwin and surrounds possessed *bimA*_Bp_ (10). Bivariate statistical analysis demonstrated that the proportion of LPS B strains carrying *bimA*_Bm_ (n = 8, 21%) is higher compared to LPS A (n = 8, 2.6%) and this was significant (P<0.001). We next determined if *bimA*_Bp_ or *bimA*_Bm_ has a similar geographical footprint to LPS A or LPS B in the Top End. (10). Geographical mapping of the *bimA* variants in the environmental strains demonstrated that *bimA*_Bp_ was linked with Darwin and surrounds (Fig 3B), having a similar geographical footprint as LPS A. In contrast, *bimA*_Bm_ was associated with the East Arnhem and the Katherine and Katherine remote region (13 of 17 (76%) environmental *bimA* strains), similar to LPS B. We also noted that the majority of the LPS B strains in the Katherine and East Arnhem region also carried *bimA*_Bm_ (n = 8/13; 62%), and these strains were concentrated in two geographical regions (Fig 3, star [n = 6/9] and triangle [n = 2/2]).

**Fig 3.**
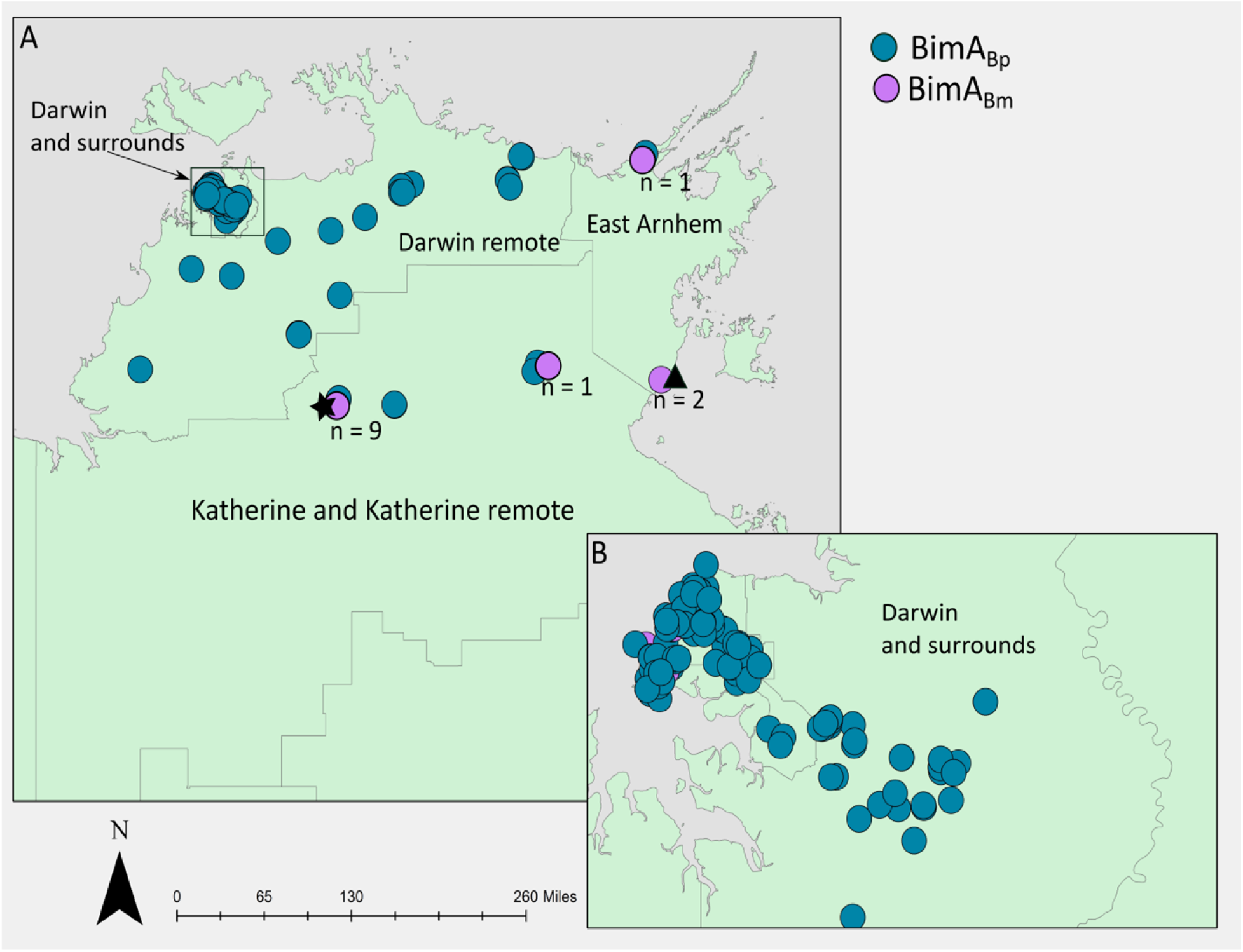
Geographical distribution of *bimA* variants in 340 environmental *Burkholderia pseudomallei* isolates from (A) the Top End region, Northern Territory, Australia and (B) the Darwin and surrounding region. Strains located in two Top End regions carried *bimA*_Bm_ and LPS B, as denoted by a black star (6/9 strains had *bimA*_Bm_ and LPS B) and a black triangle (2/2 strains had *bimA*_Bm_ and LPS B). ArcGIS (https://www.arcgis.com/index.html) was used for mapping isolates onto an Australian Bureau of Statistics shape file, of statistical area level 3 (https://www.abs.gov.au/).

### Phylogenomics revealed that LPS B2 is geographically restricted within the Top End

A phylogenomic analysis was performed to investigate the distribution of LPS types and strain relatedness on the whole-genome level (Fig 4). Overall, good clade structure based on geographical origin was observed, with strains belonging to the same ST clustering together and within the same LPS type, but with LPS A and B being dispersed throughout the phylogeny (Fig 4). However, this analysis revealed that 5/6 LPS B2 genomes of Top End origin clustered together, suggesting that LPS B2 is geographically restricted in the NT (Fig 4). Similarly, the LPS B2 strains with a Queensland, Australian origin grouped closely together.

**Fig 4:**
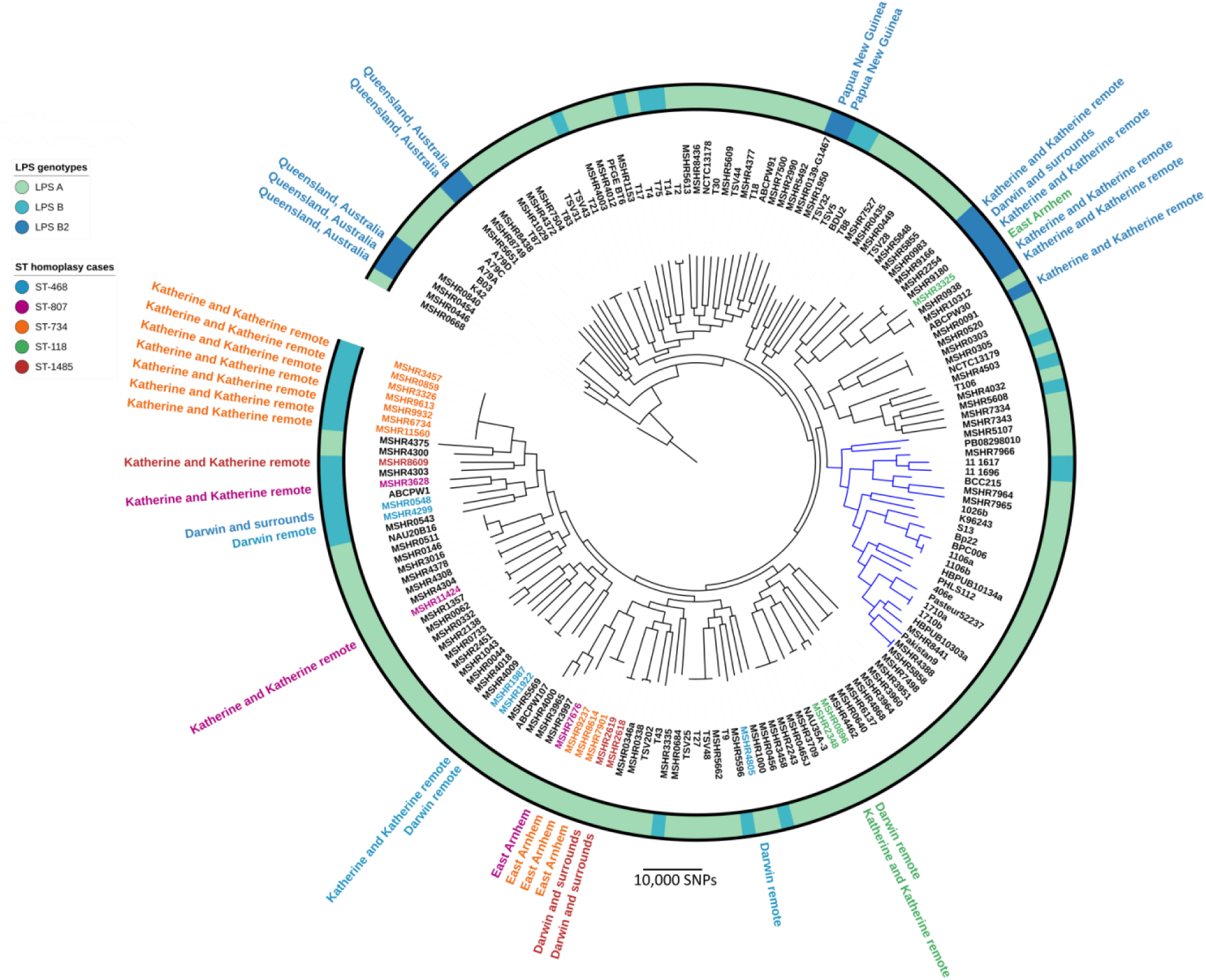
Maximum parsimony phylogeny of 175 *Burkholderia pseudomallei* genomes constructed using 219,075 biallelic SNPs. The Australian strain MSHR1153 was used as a reference for Illumina read mapping, and the tree was rooted with MSHR0668 based on the most ancestral *B. pseudomallei* isolate identified in a larger study (27). Strains with multilocus sequence type (ST) homoplasy and their corresponding geographical locations in the Top End region, Northern Territory, Australia, are indicated by the outermost text. The lipopolysaccharide (LPS) genotype of each strain is indicated by the coloured ring. Blue-coloured branches indicate the non-Australian clade. Consistency index for tree = 0.2. Interactive Tree of Life (ITOL) was used to annotate the tree (https://itol.embl.de).

### The *B. pseudomallei* LPS discrepancy in five MLST STs can be explained by ST homoplasy (isolates with the same ST that are genetically distinct)

The strong link between LPS type, geography and ST suggested that homoplasy may be present in the 18 clinical strains and three environmental strains that belonged to the five STs with more than one LPS type (Table 2). Based on comparative analysis of 175 genomes, isolates with the same ST but mixed LPS type did not group together (Fig 4: “ST homoplasy cases”) and were separated by a large number of SNPs. This is comparable to previously observed MLST homoplasy in *B. pseudomallei* (20, 21). To assess the effect of recombination on phylogenetic inference, SNP density filtering was applied to remove recombinogenic SNPs with Gubbins (v.2.3.1), which did not considerably alter the topology of the phylogenetic tree (data not shown). We also noted that the isolates belonging to ST-807 and ST-468 were from three Top End geographical regions and varied by a large number of SNPs with the largest SNP difference being ~36,000 and ~29,000 respectively (Table 2). This phenomenon represents further examples of intracontinental homoplasy but includes homoplasy occurring in more closely related geographical regions than previously described (20).

**Table 2.** *Burkholderia pseudomallei* intracontinental multilocus sequence type (ST) homoplasy cases.

| ST | Strain | Year | Sample type | Location | LPS<sup>+</sup> type | Largest SNP difference | Homoplasy present |
| --- | --- | --- | --- | --- | --- | --- | --- |
| **468** | MSHR0548 | 1998 | Clinical | Darwin and surrounds | B | ~29,000 | Yes |
|  | MSHR4299 | 2010 | Environmental<sup>#</sup> | Darwin remote | B |  |  |
|  | MSHR1987 | 2005 | Clinical | Katherine and Katherine remote | A |  |  |
|  | MSHR1922 | 2005 | Clinical | Darwin remote | A |  |  |
|  | MSHR4805 | 2011 | Clinical | Darwin remote | A |  |  |
| **807** | MSHR3628 | 2010 | Clinical | Katherine and Katherine remote | B | ~36,000 | Yes |
|  | MSHR11424 | 1998 | Clinical | Katherine and Katherine remote | A |  |  |
|  | MSHR7676 | 2012 | Clinical | East Arnhem | A |  |  |
| **734** | MSHR0859 | 1999 | Clinical | Katherine and Katherine remote | B | ~29,000 | Yes |
|  | MSHR3326 | 2008 | Clinical | Katherine and Katherine remote | B |  |  |
|  | MSHR3457 | 2009 | Clinical | Katherine and Katherine remote | B |  |  |
|  | MSHR6734 | 2012 | Clinical | Katherine and Katherine remote | B |  |  |
|  | MSHR9613 | 2016 | Environmental<sup>#</sup> | Katherine and Katherine remote | B |  |  |
|  | MSHR9932 | 2017 | Environmental<sup>#</sup> | Katherine and Katherine remote | B |  |  |
|  | MSHR11560 | 2018 | Clinical | Katherine and Katherine remote | B |  |  |
|  | MSHR7901 | 2013 | Clinical | East Arnhem | A |  |  |
|  | MSHR8614 | 2015 | Clinical | East Arnhem | A |  |  |
|  | MSHR9237 | 2016 | Clinical | East Arnhem | A |  |  |
| **118** | MSHR3325 | 2008 | Clinical | East Arnhem | B2 | ~38,000 | Yes |
|  | MSHR0896 | 1999 | Clinical | Darwin remote | A |  |  |
|  | MSHR2348 | 2006 | Clinical | Katherine and Katherine remote | A |  |  |
| **1485** | MSHR2619 | 2001 | Environmental | Darwin and surrounds | A | ~33,000 | Yes |
|  | MSHR2618 | 2001 | Environmental | Darwin and surrounds | A |  |  |
|  | MSHR8609 | 2014 | Environmental | Katherine and Katherine remote | B |  |  |

#included in study to increase ST numbers. Abbreviations: \*LPS, Lipopolysaccharide

## Discussion

Zoonotic and person-to-person transmission of *B. pseudomallei* is extremely rare, with almost every case of melioidosis therefore reflecting a single infecting exposure event from the environment in a specific geographical location. Variably present genetic markers have been documented in *B. pseudomallei* and include the mutually exclusive BimA variants (28), the BTFC/YLF gene clusters (9) and the variably present FhaB3 locus (29). The prevalence of these genetic markers varies between Australia and Asia and BTFC/YLF prevalence varies between clinical and environmental strains, and in the case of *bimA* and *fhab3* they have also been linked to clinical variation in disease phenotype (9, 10). The geographical footprint of *fhaB3* and *bimA* variants has been previously determined in the Northern Territory, with *bimA*_Bm_ associated with melioidosis in remote communities while *bimA*_Bp_ and absence of *fhaB3* is associated with melioidosis in the Darwin urban region (10).

The LPS O-antigen moiety of *B. pseudomallei* is highly varied with three serotypes described (12). Nevertheless we recently found that previously published *in vitro* differences in virulence between LPS A and LPS B did not translate to differences in mortality in DPMS patients, supporting the dominant role in human melioidosis of host risk factors in determining disease severity and outcomes (18). It has also been demonstrated that antibodies from the two dominant LPS types, LPS A and B are not cross-reactive due to structural differences of the O-antigen, which has implications for an LPS based vaccine (12). Regional differences in LPS serotype prevalence have been noted in Southeast Asia and Australia with LPS A being the most prevalent, followed by LPS B and with LPS B2 being the rarest in Australian strains and as yet undetected outside Australia (12). Despite the reporting of these regional differences in LPS prevalence, no studies have investigated the geographical footprint of the LPS types in a contiguous melioidosis-endemic region. We used a combination of statistical analysis, WGS and geographical mapping to examine the geographical footprint of the LPS genotypes in the Top End of the Northern Territory.

Our data confirm that the prevalence of the LPS types in the environmental samples (LPS A = 89%; LPS B = 10% and LPS B2 = 1%) correlates with what we previously observed in a large clinical isolate collection (LPS A = 87%; LPS B = 12% and LPS B2 = 1%) (18) and this high LPS A prevalence was also seen in the strains from Western Australia, Queensland and Papua New Guinea. High prevalence of LPS A is reflected in it also being present in many near-neighbour species (30), with the possibility of horizontal gene transfer of LPS A between these related species. LPS A was also distributed throughout the *B. pseudomallei* phylogeny being present in genetically distinct strains, also supporting horizontal transfer of the entire LPS A loci. Based on our large clinical dataset (n = 999) we found strong geographical clustering of LPS types, with the highest odds of having a clinical isolate with LPS B being in the East Arnhem region in comparison to the Darwin region (OR 19.5, 95% CI 9.1 – 42.0; *p*<0.001. A similar trend was observed for strains isolated from the Northern Territory environment (n = 340), and in the environmental strains LPS B was not detected in urban Darwin at all. We also found that LPS A was associated with STs that are highly prevalent in Darwin and surrounds (ST-109, ST-36 *p*<0.001 and ST-132). These results demonstrate that at least within the Top End context the LPS genotypes can be used as geographical indicators alongside *fhaB3* and the *bimA* variants.

Despite the strong link noted between ST and LPS type, we detected an LPS discrepancy in 21 isolates belonging to five rare STs: ST-118 (n = 3), ST-734 (n = 10), ST-468 (n = 5), ST-807 (n = 3) and ST-1485 (n = 3). Phylogenomic analysis of 175 genomes confirmed that the isolates belonging to the five STs represented five new ST homoplasy occurrences, explaining the LPS discrepancy noted in the five STs. Isolates of each ST were separated by a large number of SNPs (~29,000-~39,000), which is characteristic of Australian isolates that belong to different STs (20, 21). Despite the large number of SNPs separating isolates within each ST, they all still resided within the Australian clade, representing five novel intracontinental ST homoplasy cases. These findings further highlight the low resolution of MLST and the need for high resolution techniques such as WGS to resolve unexpected genotyping results, particularly for *B. pseudomallei* which is a highly recombinogenic pathogen (31).

Lastly, we examined the geographical footprint of the *bimA* variants using corresponding environmental strain data. Firstly, we revealed a similar geographical trend for the *bimA* variants to that previously observed in clinical strains (10). Using coordinates from the same environmental strains used for the LPS work we revealed that the rare *bimA* variant, *bimA*_Bm_, has a similar geographical footprint to LPS B, being associated with the Katherine and East Arnhem regions and the geographical footprint of the dominant *bimA* variant, *bimA*_Bp,_ is similar to LPS A being associated with Darwin and surrounds.

An association between *bimA*_Bm_ and neurological melioidosis has been described (10). Although we recently demonstrated that LPS type alone did not confer a clinical or mortality differential in human melioidosis, further studies are required to see if the combination of *bimA*_Bm_ with LPS B found in eight environmental strains from the Katherine and Katherine remote and East Arnhem regions in this study may have specific clinical implications.

The present study encompassed a number of limitations. One limitation of this study is that the assigned presumptive location of infection for cases of melioidosis is dependent on good epidemiological history and will sometimes not be the true location of infection. Nevertheless, the close correlation of geographical LPS footprints between the clinical case isolates and the environmental isolates (which are 100% location accurate) is reassuring that the prospective nature of the Darwin melioidosis study is providing accurate epidemiological data. A second limitation is that the four regions (Darwin and surrounding, Darwin remote, Katherine and Katherine remote and East Arnhem) vary substantially in both geographic size and population density, which results in a greater number of isolates obtained from the Darwin and surrounding region and ultimately clustered data. We accounted for the clustered nature of our data by using bivariate logistics regression with the cluster feature.

In conclusion, by combining genomic data with corresponding strain geographical information we have found that in the tropical north of Australia the LPS types have distinct geographical footprints as well as ST associations, adding to the already known variably present genetic markers *fhaB3* and the *bimA* variants. A novel and interesting finding was that the geographical footprint of LPS B and *bimA*_Bm_ are similar and in remote Top End locations. The clinical, sero-diagnostic and vaccine implications of geographically-based *B. pseudomallei* LPS and other gene differentials and their relationships to regional and global dispersal of melioidosis require global collaborations with further analysis of larger clinically and geospatially-linked datasets.

## Acknowledgements

We thank our microbiology laboratory colleagues at Royal Darwin Hospital for their support and expertise in *B. pseudomallei* identification. We also thank Glenda Harrington (Menzies School of Health Research) for laboratory assistance. We thank Jeffrey Warner and Anthony Baker for providing *B. pseudomallei* isolates from Queensland.

## Suppporting information Captions

**Table S1. Summary of 175 *Burkholderia pseudomallei* genomes used to construct the phylogeetic tree with LPS type, ST, origin and GenBank accession no.**

